# Alternating dynamics of segregation and integration in human brain functional networks during working-memory task

**DOI:** 10.1101/082438

**Authors:** Antonio G. Zippo, Pasquale A. Della Rosa, Isabella Castiglioni, Gabriele E. M. Biella

## Abstract

Brain functional networks show high variability in short time windows but mechanisms governing these transient dynamics still remain unknown. In this work we studied the temporal evolution of functional brain networks involved in a working memory task while recording high-density electroencephalography in human normal subjects. We found that functional brain networks showed an initial phase characterized by an increase of the functional segregation index followed by a second phase where the functional segregation fell down and the functional integration prevailed. Notably, wrong trials were associated with different sequences of the segregation-integration profile and measures of network centrality and modularity were able to catch crucial aspects of the oscillatory network dynamics. Additionally, computational investigations further supported the experimental results. The brain functional organization may respond to the information processing demand of a working memory task following a 2-step atomic scheme wherein segregation and integration alternately dominate the functional configurations.

## Introduction

The human brain can be portrayed as a giant complex network from the twofold point of view of anatomical and functional perspectives, the former probing the stable structural connections among neurons or neuronal populations, the latter focusing on the functional connections exiting in the huge dynamic repertoire of various transient outputs (actions, perceptions, cognition, etc.) (Tononi, Edelman, and Sporns 1998; Bullmore and Sporns 2009; Bullmore and Sporns 2012; Park and Friston 2013). From the functional standpoint, major efforts have been spent to provide quantitative appraisal of brain network dynamic events in different experimental or clinical conditions. Two functional states of brain networks represent a generalized hallmark of brain network dynamics: the functional segregation represented by mutual functional independence of the brain districts, and its counterpart, the functional integration, which is the ability of the brain to efficiently combine information from different regions. So far, these principia represent one of the most important paradigms in brain physiology and lay their roots in the realization that brain networks are organized in modules and in a core of densely interconnected hubs. Modules endorse the ability of brain networks to segregate information while core hubs provide the integration substrate. Many authors also reported brain topological configurations coherent with the small-world network model (Watts and Strogatz 1998) enriched by a core-periphery organization (Martijn P van den Heuvel and Sporns 2011).

So far, most studies on the brain functional connectivity have been carried out on Blood Oxygen Level Dependent (BOLD) signals in fMRI (functional magnetic resonance imaging). A drawback of fMRI signals is inherent to slowness of the BOLD signal which peaks about 2 seconds after the neural activity, collapsing the great variability of brain networks highlighted at high temporal resolutions (Whitlow, Casanova, and Maldjian 2011; Hutchison, Womelsdorf, Allen, et al. 2013; Chu et al. 2012). This coarse assumption of stationarity violates most of the brain information processing time-scales which take place over tens or hundreds of milliseconds (Park and Friston 2013; Sporns 2013a; Sporns 2013b).

Despite the brain networks intrinsically and dramatically change over time, the current knowledge about functional networks is primarily described with heavy stationarity assumptions.

The scope of this work is to study the brain topological information processes within event coherent time-scales by high density electroencephalography (EEG), able to capture neural events and to map their functional organization in due intervals, during a simple cognitive task, namely the n-back working memory test. In equivalent terms, this means to develop a model for functional brain network non-stationarity during the cognitive task. To this purpose we recorded the electroencephalographic activity of 21 healthy volunteers involved in a visual working memory task in order to elicit co-activations of large brain regions with 128 channels electroencephalogram. Indeed, previous works suggest that interareal phase synchrony sustains the object representation and the information maintenance in visual working memory tasks (Palva et al. 2013; Blinowska et al. 2013).

Results showed that during the execution of a task, brain networks encountered a first stage dominated by functional segregation followed by a second stage where functional integration prevailed. When participants failed to choose the correct answer, we observed different trends suggesting that the dynamics observed in correct trials appear not only concurrent but necessary to achieve satisfactory cognitive performances. Deeper network analyses revealed that the working load of nodes and their core-periphery organization play crucial roles in such network dynamics. Further computational investigations corroborated experimental results and built a formal explanatory theory of the discovered phenomenon.

In conclusion, this work suggests that the brain information processing, at least in working memory tasks, may occur following a 2-step hierarchical elaboration scheme where, as a rule, information is first parallelized on brain modules and subsequently integrated into core hub modules.

## Materials and Methods

### Ethical Statement

The experiment was conducted with the understanding and written consent of each participant according to the Declaration of Helsinki (BMJ 1991; 302: 1194) and in compliance with the APA ethical standards for the treatment of human volunteers (1992, American Psychological Association). The ethical committee of the Carlo Besta Neurological Institute (Milan, Italy) approved the experimental protocol. The whole experiment lasted about one hour and volunteers were not paid for their participation.

### Experiment Description and EEG Acquisition

We selected 21 young adult subjects (age average = 25, SD = 4; male = 11) and we used a freely available software implementation of the N-back working memory task (Jaeggi et al. 2003). None of the volunteers was taking psychoactive medication. Criteria for selection considered anatomical features of the head in order to fit requirements of our EEG cap (GSN-HydroCel-128, EGI). Subjects were previously instructed about the graphical task interface and a short toy session of the 1-back task was allowed to get a good familiarity with the user interface. Subjects underwent 3 sessions of 41 trials of 1-back task and 3 sessions of 41 trials of 2-back task. The task was constituted by a sequential presentation of square-shaped boxes, which appeared on the screen. Subjects had to keep in mind the color and the position on the screen (out of nine possible). In each trial a colored box appeared on a randomly selected position of the grid for 500 ms and the subject was asked to respond within 2500 ms by possibly pressing 1 of 2 buttons to indicate whether there was a color or a position match (or both) between the current box and the one seen in the previous (one previous for 1-back and two previous for 2-back) trials. Since the used N-back software did not track the timestamps of user responses, we redirected the input system into a Sony Playstation GamePad whose buttons were covered by touch sensors that delivered signals both to the N-back software and to the EGI amplifier.

The subjects were comfortably seated, their arms leaning on a surface to avoid muscle contraction interference and their feet placed on a platform. Participants performed variably on the working-memory tasks averaged and performance was generally good. Mean percentage of correct trials was 94.1% ± 6.8 (SD) for 1-back sessions and 89.3% ± 10.6 (SD) for 2-back sessions; mean reaction time was 0.56s ± 0.19.

We recorded electroencephalographic activity with a EGI Net Station 400 equipped with a 128 electrodes GSN-HydroCel cap. The cap was positioned according to the vendor guidelines by matching three reference electrodes around the scalp. Prior to acquisition, we measured amplifier gains and electrode impedances. We fixed electrodes with impedance values greater than 70 KOhm by adding few drops of a hydrosaline solution between the electrode sponge and the scalp. We followed this procedure until all electrode impedances were below the threshold. Electroencephalographic signals were acquired at a sampling frequency of 500 Hz. The whole recording lasted for around 30 minutes.

Since we had no tool to measure the exact position of electrodes or to elucidate the anatomical substrate of participants (e.g. MRI) we excluded any further investigation that involved reliable structural information (cortical mapping, source localization, etc.).

### EEG Processing

EEG recorded sessions were processed in Matlab with the *eeglab* toolbox^34^ and with “in house” developed routines. Raw signals were mean corrected and filtered in the frequency range [12,40] Hz of interest for beta and gamma bands. Additionally, signals were filtered to attenuate line noise at 50 Hz using a 0.3 Hz width notch filter. To remove physiological (eye movement, respiration, heartbeat) and extraphysiological (instrument, environment) artifacts, we performed an Independent Component Analysis (ICA) (Delorme and Makeig 2004) of the signals using the standard algorithm provided in the eeglab toolbox (*runica*). A meticulous visual inspection classified bad independent components that were thus removed from the EEG signals.

Subsequently EEG signals were split into 246 epochs corresponding to the 41 trials of 6 n-back sessions. We considered the start of each trial 200 milliseconds before the visual presentation of the box on the screen grid and the end 2800 ms after such event obtaining 10 time windows for each trial. We considered different sizes variable from 500 ms to 2 seconds to establish possible conditionings but the obtained networks and their related statistics did not significantly change. Indeed, the chosen time window length was short enough to capture great variations of functional connections though preserving robustness of the estimated synchronization index (WPLI).

### Functional Connection Extraction

The scope of this work was to investigate the fast functional network changes occurring in a cognitive task. To extract the functional connections among electrodes in each trial, we evaluated several methods based on synchronization and after a throughout evaluation, we chose the weighted phase lag index (WPLI) because it is capable to minimize effects of volume conduction which tightly affect high density EEG recordings (Vinck, Oostenveld, van Wingerden, et al. 2011; Gordon et al. 2013).

Formally, let *Z* and *iZ* are respectively the real and imaginary parts of the cross-spectrum of two EEG signals x and y. The Weighted Phase-Lag Index can be defined as:

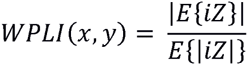

where E is the expected value and |⋅| is the absolute value function. The inequality 0 ≤ *WPLI*(*x*, *y*) ≤ 1 holds for each couple of signals *x* and *y* and *WPLI*(*x*, *y*) = 1 if *x* and *y* are maximally synchronized while *WPLI*(*x*, *y*) = 0 when there is no synchronization at all. Functional connectivity graphs were represented by adjacency matrices A obtained by computing *A*(*i*, *j*) = *WPLI*(*i*, *j*) for each (*i*, *j*) couple of EEG electrodes (*i*, *j* ∈ {1, …, 128}).

### Network Analysis and Comparison

For the analysis of these graphs, we introduced a set of common statistics from the Complex Network Theory able to estimate the network extent to integrate and segregate information and many other relevant network features (see Table 1). Functional segregation and integration in networks can be measured by two statistics: the clustering coefficient (C) and the characteristic path length (L). The former measures how close the neighbors of a node are to being a clique. The latter estimates the average shortest path length in the graph, i.e. how much the nodes are accessible. Computationally, analyses on the extracted functional brain networks were performed in Matlab by the Brain Connectivity Toolbox (BCT) and by other routines developed in our lab (Rubinov and Sporns 2010).

**Table 1:** The complex network statistics used in this work. We reported the weighted versions of each statistic. Weights are assumed to span from 0 to 1.

| Measure | Definition | Interpretation |
| --- | --- | --- |
| Node strength | $k_i = \sum_{j \in N} a_{ij}$ | Number of edges connected to a given node $i$. Nodes with relatively high values of $k$ are called *hubs* |
| Shortest path length | $d_{ij} = \sum_{a_{fg} \in r_{i \leftrightarrow j}} a_{fg}$<br>where $r_{i \leftrightarrow j}$ is the shortest path between $i$ and $j$ | The number of edges encountered in the shortest path between node $i$ and $j$ |
| Characteristic path length | $L = \frac{1}{n} \sum_{i \in N} L_i = \frac{1}{n} \sum_{i \in N} \frac{\sum_{j \in N, j \neq i} d_{ij}}{n-1}$ | Measure of network integration |
| Clustering coefficient | $C = \frac{1}{n} \sum_{i \in N} C_i = \frac{1}{n} \sum_{i \in N} \frac{2t_i}{k_i(k_i-1)},$<br>with $t_i = \frac{1}{2} \sum_{j,h \in N} a_{ij} a_{ih} a_{jh}$ | Measure of fine-grain network segregation. It counts the average number of triangles $t$ (3-node fully connected graphs) present in the network |
| Modularity | $Q = \frac{1}{l} \sum_{u,v \in N} \left[ a_{uv} - \frac{\Omega_i \Omega_j}{l} \right] \delta_{m_i} \delta_{m_j},$<br>where $l$ is the sum of all weights of $V$ (whose elements are called *modules*) and $m_i$ is the module containing the node $i$ and $\delta_{m_i} \delta_{m_j} = 1$ if $m_i = m_j$ and 0 otherwise. | It evaluates the tendency of the network to be reduced in independent (or scarcely dependent) modules |
| Eigenvector Centrality | $EC_i = \frac{1}{\lambda} \sum_{z \in Z(v)} x_z,$<br>where $Z(v)$ is a set of neighbors of $v$ and $\lambda$ is a constant | It assigns relative scores to nodes whose connections to high-scoring nodes contribute more to the score of the node in question than equal connections to low-scoring nodes |
| Betweenness Centrality | $BC_i = \frac{1}{(n-1)(n-2)} \sum_{h,j \in N, h \neq j, h \neq i, i \neq j} \frac{\rho_{hj}(i)}{\rho_{hj}},$<br>where $\rho_{hj}$ is the number of shortest paths between $h$ and $j$, and $\rho_{hj}(i)$ is the number of shortest paths between $h$ and $j$ that pass through $i$ | It is the amount of shortest paths that pass through the node $i$. It roughly indicates how much information burdens the node $i$ |
| Small-worldness | $S = \frac{C/C^r}{L/L^r}$ | The indices quantify the affinity of a network to be a small-world network. $S$ should be greater than 1 and $w$ close to 0 |
| | $\omega = \frac{L^r}{L} - \frac{C}{C^l}$ | |

In complex network theory, several graph measures take specific meaning only if they are compared to the same graphs subjected either to randomization or to latticization (often called *null networks*). Both procedures guarantee that the node degree distributions of the original graphs were preserved. We computed, by using the Matlab function *randmio_und.m*, the randomized version of our graphs and we estimated *C*^*r*^ and *L*^*r*^ (*C*^*l*^ by latticization, *latmio_und.m*). These null network values are required to verify the small-world nature of the graphs. In fact, classical and novel measures of small-worldness such as *S* and ω state that, respectively, if S > 1 and ω (which takes values in [−1,1]) is close to 0, the graph can be considered a small-world network (Stam, Nolte, and Daffertshofer 2007; Humphries and Gurney 2008).

The functional graphs obtained by our analysis were further characterized to study the information workflow. To this aim, we estimated the network centrality with the notions of betweenness and eigenvector centrality (Borgatti 2005; Gould 2016). Because it can be interpreted as a measure of the importance of a node within the network, the distribution of node centrality highlights how the information workflow is distributed among nodes.

We further studied the modularity and the community structure of our graphs but preliminary analyses performed with the two most used community detection algorithms (Freeman 1977; Telesford et al. 2011) reported modularity values close to zero nullifying the validity of the computed partitions. The problem was due to the well-known tendency of such algorithms to prefer clusters of big size. Therefore, we decided to tweak the multi-resolution parameter in order to force the Louvain algorithm to produce modules of smaller dimension (gamma = 1.15 in all analyses). By using this trick, we obtained acceptable values of modularity (Q > 0.15 for all partitions) and proceeded with inferences.

Ultimately, we analyzed networks that evolved in time potentially dropping and recruiting nodes and connections that come from different experimental conditions. Such a methodology requires the discussion of potential issues. We performed network analyses on the original weighted version of graphs instead of using binarization techniques for several reasons. First, unconnected nodes can be rare but could occur especially after adjacency matrices binarization. Network comparisons with different number of nodes require hard statistical analyses and to discard incompatible networks (for instance those with too few nodes or with more than one strongly connected component) from observations. Second, graph thresholding produces inevitably loss of information. Third, the functional connectivity measure (WPLI) has also been proposed to enrich the reduced dynamic range conveyed by its previous version (the PLI) especially for weak interactions (Vinck, Oostenveld, van Wingerden, et al. 2011). Hence the selective removal of portions of weights almost surely will reduce the power of the consequent network statistics. Four, when thresholding graphs, it is necessary to repeat computations for a discrete range of subjective values thus producing a considerable increase of the overall computational complexity (see Discussion). Last, all network statistics that we needed for our analyses have a weighted counterpart (Rubinov and Sporns 2011).

### Computational Network Models

Computational models are commonly used to investigate complex system dynamics. In this study, we tried to figure out which factors led the observed topological dynamics by simulating information flow within network models coherent with the current knowledge of the brain topologies. Thus we produced two groups of network models, one that contains two network models consistent with brain topologies and the other one contains two null networks as hypotheses to be rejected. Specifically, we considered the Watts-Strogatz model (WS) and the Barabasi-Albert model (BA) to generate, respectively, small-world and core-periphery networks (van den Heuvel et al. 2008). On the other hand, we examined Erdős-Renyi (ER) and the ring-lattice (RL) models to generate totally random and purely deterministic networks representing the null hypotheses.

The information flow within networks was simulated starting from the notion of edge betweenness centrality (EBC, closely related to betweenness centrality) which assigns at each edge a centrality score in a similar manner of BC, namely by counting the number of times each edges is involved in a shortest path between all node couples.

Although the alternating behavior observed in the experimental data was evolved in 10 time windows, the core dynamic was limited in two phases, therefore we divided the time of flow in two parts within we expected to observe the working hypothesis.

In this edge-centric perspective, to simulate the network information flow within the two time buckets we computed the rank statistics of the EBC edge distribution which allowed us to arbitrary part the edges involved in the first phase from those recruited in the second one. Since we could not assume the number of activated edges in each phase, we inspected all possible combination with a resolution of one percentile. Therefore, we computed and collected both functional graphs for each of the 100 percentiles that served as leverage points along the EBC distribution. Essentially, by keeping a certain set of edges in the first phase we considered those edges as activated and thus belonged to the same functional graph for the first time bucket. The complementary set of edges not activated in phase 1 fell in the time bucket 2 assembling another functional graph. For both functional graphs we computed values of functional integration (L) and segregation (C) and each of the 4 network model was exerted 100 times to reduce effects of randomness. As a rule, for each generated network we computed the same network statistics (C and L) of the entire graph (all edges together) to track the magnitude of functional evoked network modifications because this network represented the structural substrate of the evoked functional graphs.

In addition, to elude possible effect of the network dimension, we investigated such dynamics for variable network sizes spanning 6 dyadic scales (2^5^ to 2^10^, higher scales were prohibitively costly in terms of computational time) and for each network model. The entire computational framework is freely available, and it is able to reproduce data and figures (https://sites.google.com/site/antoniogiulianozippo/codes, section “Information Flow in Brain Network Topologies”).

### Statistical Tests

In this study we used different type of statistical hypothesis tests. When we had to assert the significance of the Pearson’s correlation coefficient, we used the permutation test by performing 10000 permutation of the dataset and evaluating the number of times (out of 10000) the computed correlation was greater or equal of that computed on the original dataset. Whether the number of times was smaller than 500 (equivalent to get p < 0.05) the null hypothesis was rejected.

When we had to assess if two samples come from different distributions we used the non-parametric two-paired Wilcoxon ranksum test. Similarly, in case of more than two samples, we used the Kruskal-Wallis test. We preferred non-parametric tests because we had no confidence about the normal distribution of the samples. Eventually, when we directly compared two distributions we used the Kolmogorov-Smirnov test.

Within the Results section, “P” indicates the level of significance and “N” indicates the sample dimensions. Statistical quantities are reported by using three decimal digits.

## Results

### Functional Network Features

We investigated the dynamics of brain functional networks in a simple working memory task with high density electroencephalographic recordings in young healthy adults. The working memory (WM) is responsible for the storage and maintenance enabling integration of higher order information (Baddeley and Wilson 2002; Baddeley 2003; K Oberauer and Suss 2000; Klaus Oberauer 2003; Pessoa et al. 2002). WM capacity is expressed through a high level of stability across different cognitive functions, thus relying upon the organization and interaction of multiple brain regions ruled by dynamic changes in cognitive control systems(Jaeggi et al. 2003; Owen et al. 2005).

In this study, we were interested in unraveling short-term dynamics, in terms of brain functional connectivity features, while subjects performed the n-back task, and to trace the information flow within WM functional networks. In the n-back task, participants are required to monitor a stream of visual stimuli (colored squared boxes), presented one at a time. The task is to indicate whether the currently displayed item is identical as the one presented n trials previously. The memory demands grow as n increases. In the present study, n was set at 1 or 2.

We recorded ten minutes of ongoing (resting) activity before the beginning of the cognitive task and six sequences of 41 trials (i.e. 3 sequences of 1-back and 3 sequences of 2-back). Visual patterns appeared on the screen for 0.5 seconds at 3 seconds intervals. In order to study the non-stationarity, we split each recorded trial into 10 sliding time windows, 1 second width each with a sliding step of 200 ms (see Materials and Methods and Table 2). To extract the functional connections among the 128 electrodes, we evaluated several methods based on synchronization choosing the weighted phase-lag index (WPLI, Vinck, Oostenveld, Van Wingerden, et al. 2011; Ortiz et al. 2012) thus minimizing the effects of volume conduction that strongly affects EEG recordings. We selected the frequency bands of beta and low gamma [12-40 Hz] where we found conspicuous transients of evoked potentials along the temporal, fronto and parietal cortical areas (Figure 1E-F).

**Figure 1.**
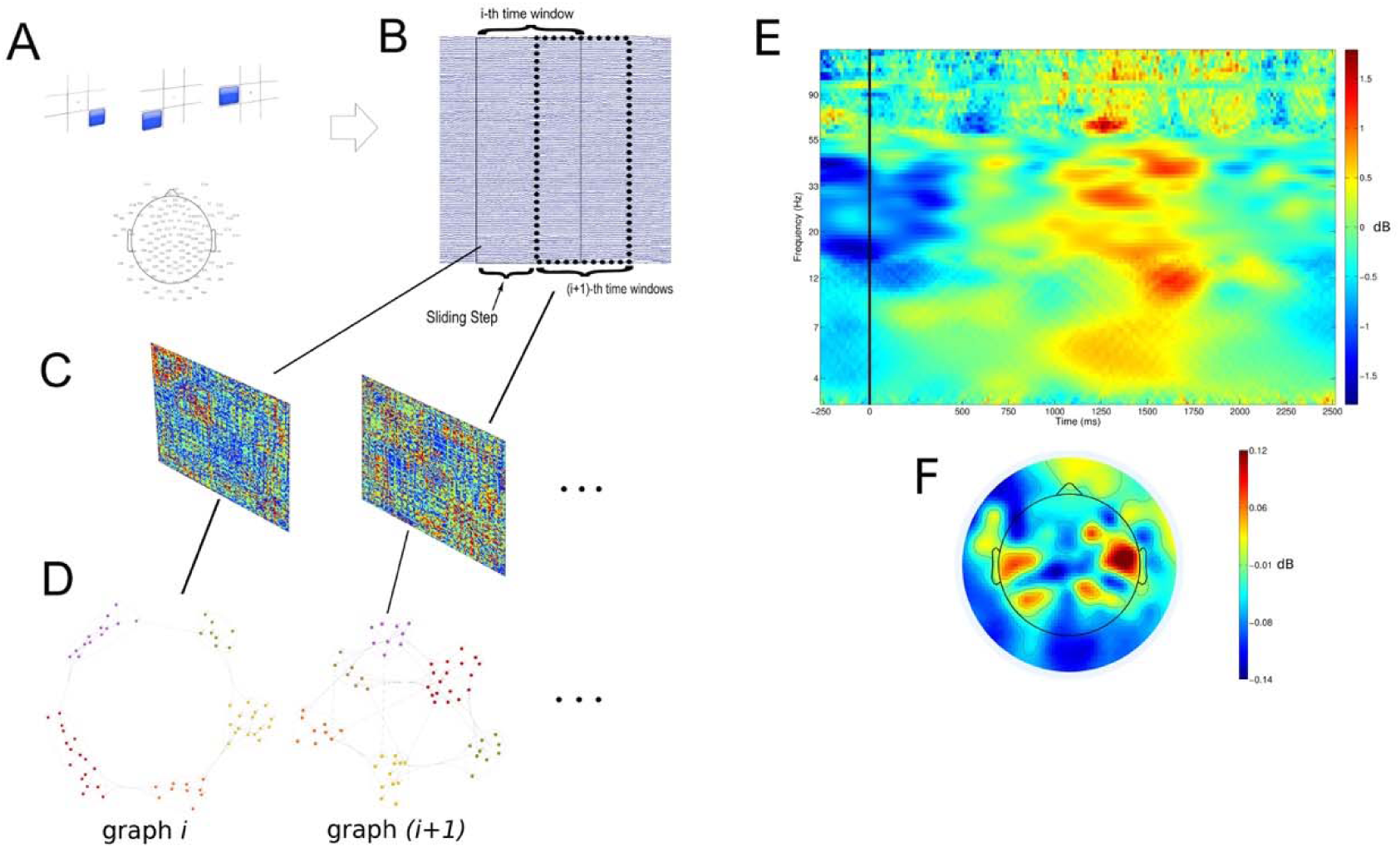
Design of the experimental framework. (A) Electrode locations in a two-dimensional mapping of human scalp. Locations are referred to the standard positions of the GSN-HydroCel-128 EGI cap in the BESA sphere space. (B) Example of the windowing mechanism used in the study. The i^th^ window is followed by the partially overlapped (i+1)^th^ window. EEG signals from each window produce the extraction of an adjacency matrix (C) by computing the WPLI value for each couple of EEG signals. (D) Graphs obtained from the above adjacency matrices where nodes are displayed according to a community layout. (E) Average Evoked Potentials among all subjects and trials, the most powerful and lasting frequency bands elicited by trials were the beta and low gamma (12-40 Hz) mainly distributed over the parietal and mediotemporal lobes (F).

**Table 2:** Time windows in task analysis. All windows had a width of 1000 ms and the sliding step was of 200 ms. Each trial lasted 3 seconds and for analysis we extracted data samples −200 ms before and 2600 ms after the presentation of the visual pattern.

| Window number | Left bound | Right bound | Centroid |
| --- | --- | --- | --- |
| 1 | -200 ms | 800 ms | 300 ms |
| 2 | 0 ms | 1000 ms | 500 ms |
| 3 | 200 ms | 1200 ms | 700 ms |
| 4 | 400 ms | 1400 ms | 900 ms |
| 5 | 600 ms | 1600 ms | 1100 ms |
| 6 | 800 ms | 1800 ms | 1300 ms |
| 7 | 1000 ms | 2000 ms | 1500 ms |
| 8 | 1200 ms | 2200 ms | 1700 ms |
| 9 | 1400 ms | 2400 ms | 1900 ms |
| 10 | 1600 ms | 2600 ms | 2100 ms |

We analyzed the functional connectivity graphs with a set of common network statistics preferring to keep graphs in their original weighted form(Rubinov and Sporns 2010). The extent of functional segregation was estimated by means of two graph-related measures, namely the clustering coefficient (C) and the functional integration through the characteristic path length (L, Tononi, Edelman, and Sporns 1998; Rubinov and Sporns 2010). To investigate the behavior of functional segregation and integration in different experimental conditions we collected networks statistics from all recording windows and proceeded with analyses and inferences (Figure 1A-D reports the analytic approach). The columns of Table 3 summarize basic network statistics for functional connectivity graphs obtained in 3 different conditions: resting state, 1-back and 2-back tasks. Specifically, we found a tight general correlation between C and L (R = 0.8973, P = 0.000, permutation test, N = 33600). Subsequently we analyzed C and L in the different experimental conditions and we found that the cognitive task (“Overall Trials” in Table 3) produced an increment of C (P = 0.007, N = 33600, non-parametric Wilcoxon ranksum test) and a decrement of L (P = 0.014, N = 33600, ranksum test). Furthermore, C values in 1-back trials were smaller than C in 2-back trials (P = 0.000, N = 33600, ranksum test) and L values in 1-back trials were greater than in L in 2-back trials (P = 0.000, N = 33600, ranksum test). These analyses evidenced that the cognitive tasks produced significant changes in the functional networks and that the task difficulty was proportional to the extent of functional segregation and integration.

**Table 3:**
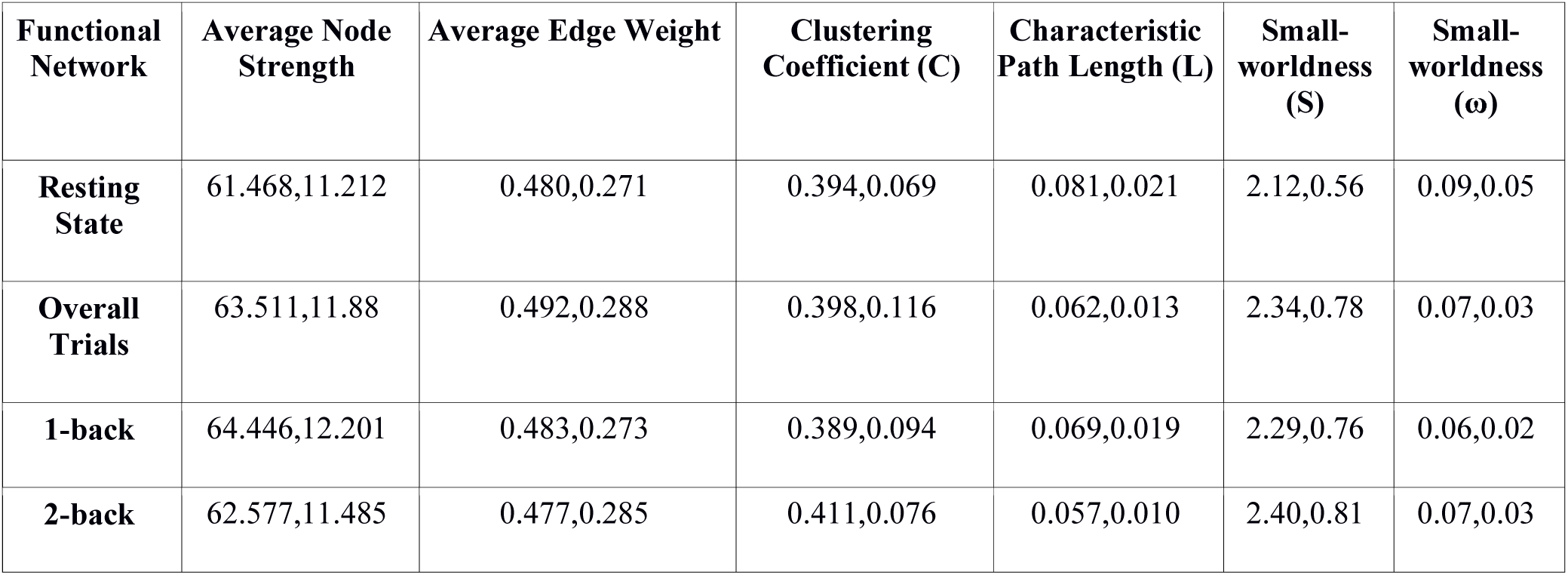
Functional brain network general statistics (mean value, standard deviation). All networks had 128 nodes.

| Functional Network | Average Node Strength | Average Edge Weight | Clustering Coefficient (C) | Characteristic Path Length (L) | Small-worldness (S) | Small-worldness ($\omega$) |
| --- | --- | --- | --- | --- | --- | --- |
| **Resting State** | 61.468,11.212 | 0.480,0.271 | 0.394,0.069 | 0.081,0.021 | 2.12,0.56 | 0.09,0.05 |
| **Overall Trials** | 63.511,11.88 | 0.492,0.288 | 0.398,0.116 | 0.062,0.013 | 2.34,0.78 | 0.07,0.03 |
| **1-back** | 64.446,12.201 | 0.483,0.273 | 0.389,0.094 | 0.069,0.019 | 2.29,0.76 | 0.06,0.02 |
| **2-back** | 62.577,11.485 | 0.477,0.285 | 0.411,0.076 | 0.057,0.010 | 2.40,0.81 | 0.07,0.03 |

### Dynamics of Functional Segregation and Integration

We proceeded investigating the dynamics of functional segregation and integration within the time windows. By averaging the functional graph statistics on all trials for each subject (see Figure 2A-F), we found a specific trend within each n-back trial, in particular, between 300 and 700 milliseconds (time windows 2, 3, 4) after the presentation of the visual pattern, when the C value (measure of functional segregation) reached its peak then decreasing between 1300 and 1700 milliseconds (time windows 6, 7, 8), the period where, conversely, L reached its minimum (indicating, being an inverse measure of integration, a peak of functional integration). Statistical tests (Kruskal-Wallis) indicated that C and L variations were significant both considering all trials (C: P = 0.002, L: P = 0.000, N = 51660) and separately, the 1-back trials (C: P = 0.000, L: P = 0.000, N = 25830) and the 2-back trials (C: P = 0.000, L: P = 0.000, N = 25830). Times of maxima and minima were slightly variable among subjects. Again, in all trials C and L were tightly correlated (R = 0.8910, P = 0.009, permutation test). Taking into account the C and L minima and maxima in 1-back and 2-back conditions, we found that C maxima were larger (P = 0.000, N = 5166, ranksum test) and C minima were smaller (P = 0.003, N = 5166, ranksum test) in 2-back than in 1-back trials. Complementarily, L maxima were larger (P = 0.000, N = 5166, ranksum test) and L minima were larger (P=0.020, N = 5166, ranksum test) in 1-back trials. Equivalently, widths of C maximal oscillations were greater in 2-back than in 1-back while widths of L maximal oscillations were greater in 1-back.

**Figure 2.**
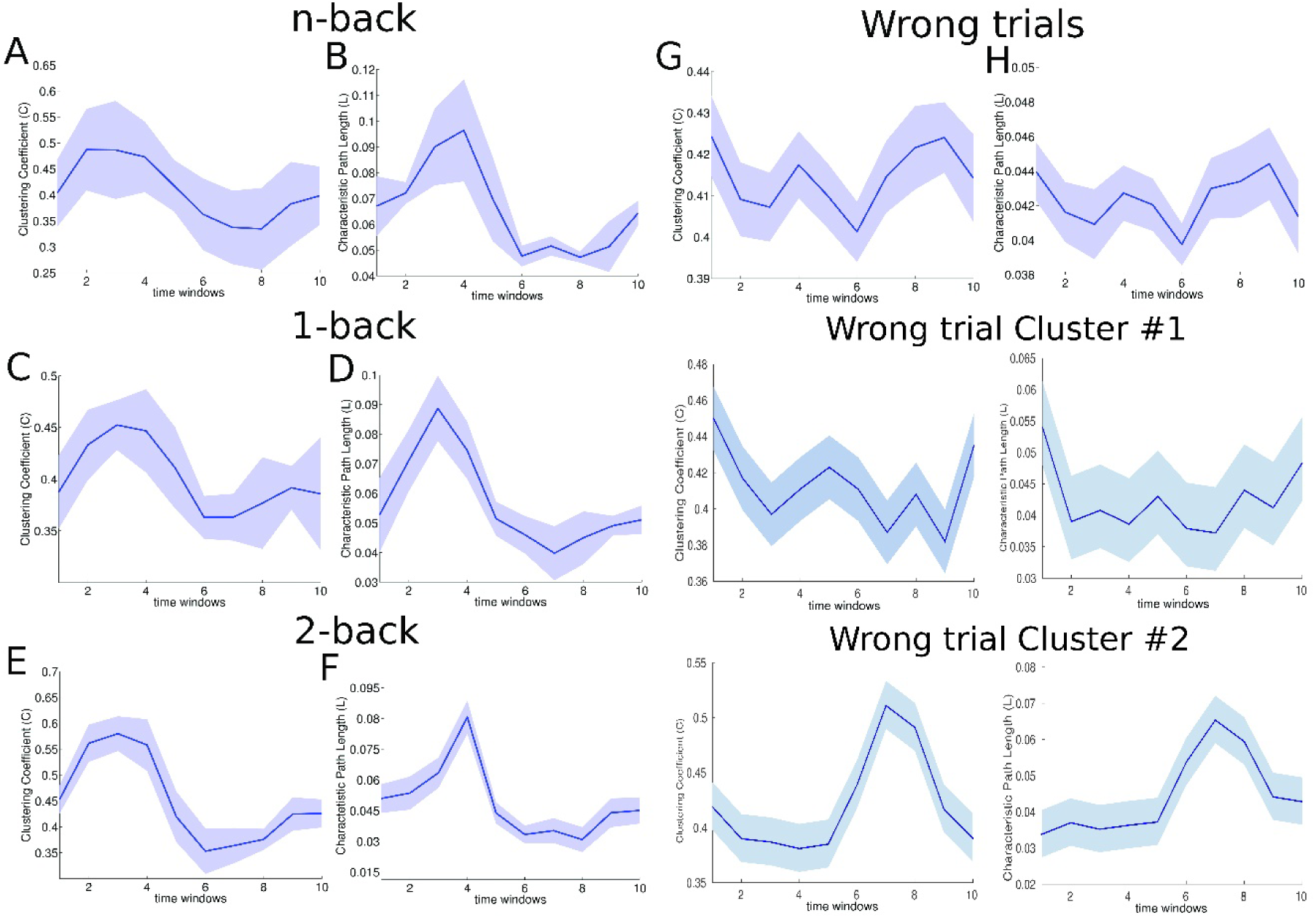
Dynamics of network statistics during the cognitive task. Lines are obtained averaging all cognitive task trials and all participants. (A-B) Figures clearly indicate a first phase spanning time windows from 2 to 4 where C and L values reached a relative maximum. The figures in (C-D) represent same statistics computed by considering only 1-back trials and in (E-F) only 2-back trials. The used windowing technique is illustrated in Figure 1 and time windows characterized in Table 2. (G-H) Trends obtained averaging all cognitive task wrong trials in all participants. Visual inspections revealed two common behaviors in C and L statistics. (I-L) The clustering coefficient was not statistical significant within the wrong trials as well as the characteristic path length. The second one (M-N) instead showed a significant modulation of the network statistics within the trials but the C and L maxima were got too late, around the seventh time window. Shaded regions indicate the standard errors.

To rule out that our results were influenced by a modulatory effect of the functional connectivity measure, we further analyzed the dynamics of the WPLI synchronization index during the task execution and we explored potential correlations with the network statistics above. We noted no significant variations of the average synchronization index along time windows (P = 0.816, N = 33600, Kruskal-Wallis test). Consequently, C and L values were not correlated to the average WPLI values (C: R = 0.1032, P = 0.023, L: R = 0.0789, P = 0.007, permutation tests, N = 33600).

Altogether these results indicate that the architecture of functional connectivity graphs shuttled between two specific states, more segregated in the first phase supervened by integration in the second. Secondarily, the results show that greater cognitive loads, as in 2-back compared with 1-back, made networks more integrated and more segregated. Finally, the observed effects were not due to modulations of the WPLI synchronization index.

Occasionally, though the percentage of correctly completed trials was high (~92% on average), participants got wrong answers to trials. In these cases, the C average distribution strongly differed from that obtained in the correct trials (P = 0.003, N = 10, Kolmogorov-Smirnov test), where the C variations within the task were not significant (P = 0.997, N = 5682, Kruskal-Wallis test). Similarly, the L average distribution was different (P = 0.001, N = 10, Kolmogorov-Smirnov test) and L variations were not significant (P = 0.914, N = 5682, Kruskal-Wallis test). Results are shown in Figure 2G-H. By analyzing the C and L behaviors in mistaken trials, we isolated two common trends depicted in Figures 2I-N. In the first kind of trend (Figure 2I-L) no modulation of C or L appeared while in the second one (Figure 2M-N) a significant modulation occurred only in the time windows 6-9. These results indicate that, when participants got wrong answers, the corresponding functional connectivity networks did not modulate integration and segregation during those trials. Notably, in the presence of delayed modulations (increments of C or L), there were associated wrong trials. Essentially, we deduced that the first 4-5 time windows correspond to the convenient interval where the first stage of network modifications (increases of functional segregation or decreases of functional integration) need to occur under penalty of a wrong trial.

### Network Centrality and Modularity during Working Memory Trials

In a second stage, we aimed to characterize functional connectivity graphs with more complex network statistics (centrality and modularity) able to highlight the information flow dynamics within networks. Typically, this requires measures that estimate the importance of nodes in the network information routing usually addressed by specialized network measures (centrality) of two types: those that use only information of neighbors (local centrality) and those that instead use information from the entire network (global centrality).

Among the local centrality measures we preferred the node degree centrality, a quantitative measure of the number of node connections, whereas, for the measures of global centrality, we chose the betweenness (BC) and the eigenvector centralities (EC, Borgatti 2005; Gould 2016). The former measures the number of times a node is bridging neighboring or far nodes. The latter assigns a greater centrality to a preeminent node (a richly connected node) than to a poorly connected one.

We found that the average node degree was weakly correlated with both C (R = 0.640, P = 0.000, N = 51660, permutation test) and L (R = 0.579, P = 0.000, N = 51660, permutation test), and that it was modulated during task trials (P = 0.007, N = 51660, Kruskal-Wallis test). Conversely, the centrality measures appeared to be invariant during task trials (BC: P = 0.999, EC: P = 0.997, N = 51660, Kruskal-Wallis tests, Figure 3A-B). This last result fostered us to further investigate the role of centrality in such brain functional networks by analyzing the distribution of the centrality measures irrespective of the trial time windows. We found that BC and EC showed heavy-tailed distributions (Figure 3C). The degree centrality on the contrary appeared normally distributed with a slight positive skewness (0.11). These results suggested that node loads were inhomogeneously distributed among nodes identifying groups of nodes that likely process a much higher amount of information than other ones. In conclusion, global centrality appears to be an important indicator of the node role during the n-back cognitive task because it identifies a stable node configuration during task completion.

**Figure 3.**
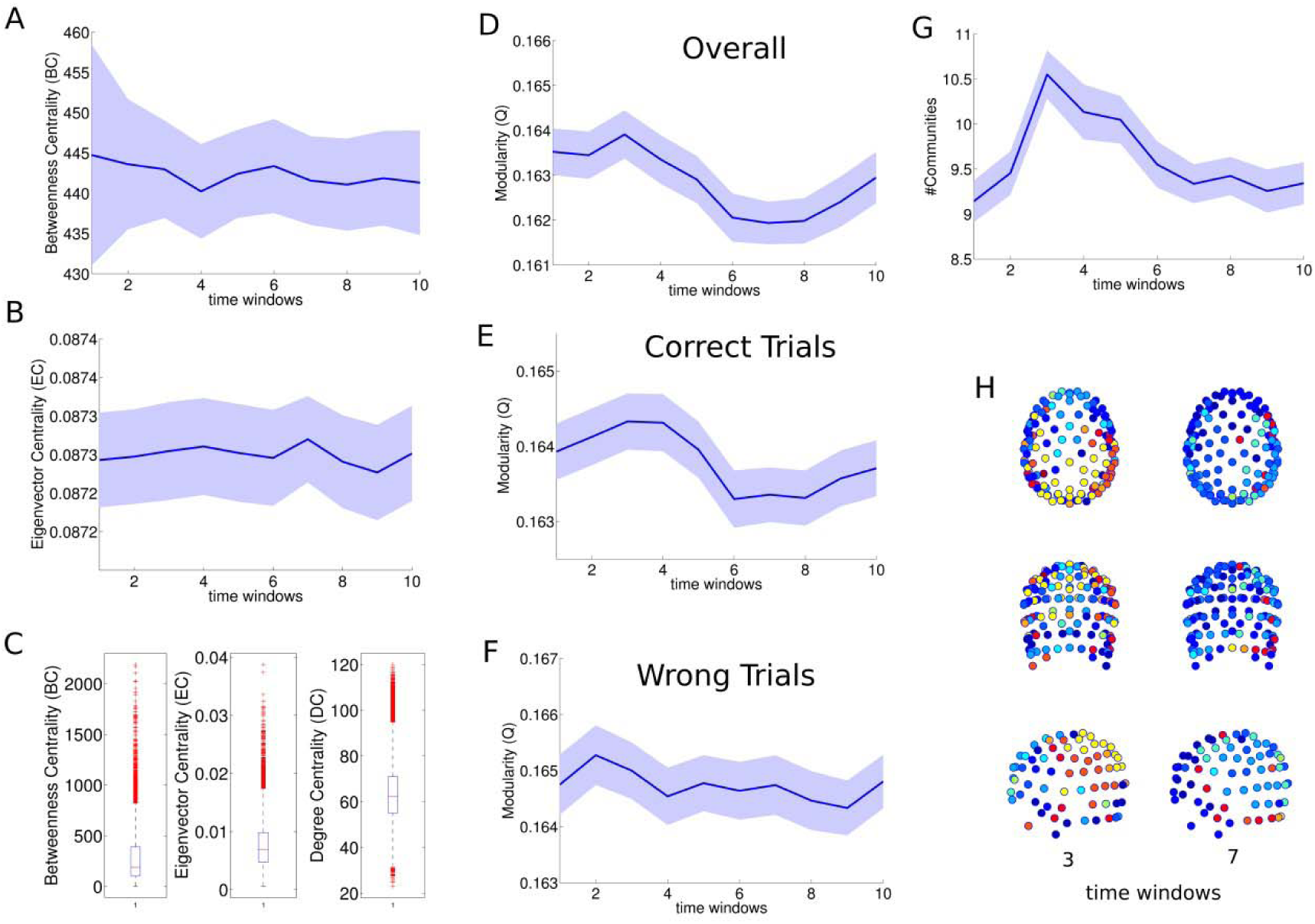
Centrality and Modularity in brain functional networks. Trends of betweenness (A) and of eigenvector (B) centralities obtained averaging all cognitive task trials and all participants. (C) Distributions of betweenness and eigenvector centralities have heavy-tailed distributions while degree centrality has a Gaussian-like shape. (D) Estimated modularity index, computed considering all trials, where is visible a straightforward dynamical effect not, however, statistical significant. (E) The modularity obtained considering only trials in which participants did correct answers, gave significant effects, with peaks of Q in time windows 2-4 and minima of Q in windows 6-8. For this reason, Q was tightly correlated with C and L. (F) Wrong trials showed no significant modulations of the modularity index. (G) The modification of the number of communities within task trials was significant: when in functional networks prevailed on the functional segregation (Figure 2), the number of communities increased of 20-30%. (H) Example of community dynamics where the left column plots represent network configurations in the time window 3 and right column plots of time window 7 of a representative trial. Shaded regions indicate the standard errors.

Since, other important aspects of complex network dynamics can be hidden in the brain modularity structure, we subsequently performed a network modularity analysis. As a rule, nodes of networks can be clustered into groups (communities or modules) in order to maximize the number of edges within each community and to minimize the number of intercommunity edges (Freeman 1977). The modularity index Q represents the goodness of the proposed partition and takes values within [0,1]. The modularity was correlated (R = 0.837, P = 0.000, N = 55166, permutation test) with the clustering coefficient indicating that the modularity increased during peaks of functional segregation and, vice versa, it declined during peaks of functional integration (R = 0.890, P = 0.009, N = 55166, permutation test). Furthermore, we found that the modularity did not significantly changed during the task trials (P = 0.322, N = 51660, Kruskal-Wallis test, Figure 3D) but when correct trials were extracted, modularity modulations became significant (P = 0.032, N = 47683, Kruskal-Wallis test, Figure 3E). Differently, this did not happen in wrong trials (P = 0.950, N = 3977, Kruskal-Wallis test, Figure 3F). Communities represent also a coarse-grain measure of the network information segregation, coupled to the clustering coefficient. Thus, by analyzing the number of communities in task trials, we found that the number of communities was significantly modulated (P = 0.016, N = 51660, Kruskal-Wallis test, Figure 3G) showing a positive slope (+20%) in the time windows 2, 3 and 4. This fact suggested that when networks were in functional segregation modality, they have more modules whereas functional integration did not require specific adjustments of the community cardinalities.

In Figure 3H, network node colors express the membership to community and in the first column (left) appear the communities in the time window 3 and in the second column (right) the communities of time window 7. Figures highlight that in earliest network configurations dominated by functional segregation preeminence, there are more communities than in the later ones where functional integration prevails. Therefore, modularity changed observing a comparable scheme to that found with a finer-grain measure of segregation (C).

### Computational Inspections

In the last stage, we tried to figure out the possible factors which enliven the alternating behavior of segregation and integration, the working hypothesis of this section. Therefore, we conjectured that some of the information processing in human brain networks could be mirrored in the observed alternating dynamics.

Hence, we developed a computational model which simulated the information flow within brain networks in comparable conditions, in order to replicate the topological dynamics observed in experimental data. Connectomic studies described the human brain as a small-world network with a strong core-periphery and scale-free organization(Newman 2006). The core-periphery feature identifies bipartite networks with a partition characterized by densely interconnected nodes with high centrality, and a complementary partition with sparsely interconnected and (usually) non-central nodes. Accordingly to these facts, since functional brain networks have both a small-world and a core-periphery organization (see Table 3 and Results sections), we challenged the working hypothesis against 4 network models: two of them coherent with brain network topologies, namely the Watts-Strogatz model (WS), able to generate small-world networks and the Barabasi-Albert model (BA), able to generate (scale-free) core-periphery networks(M P van den Heuvel et al. 2008), while on the other hand, 2 null networks, the Erdős-Renyi (ER) and the ring-lattice (RL) models, respectively a network were edges are completely randomly assigned and, conversely, a network with a purely deterministic allocation of edges.

Network dynamics can be conveniently analyzed from an edge-centric rather than a node-centric perspective following an approach similar to works of Grady et al.(Senden et al. 2014) and Ekman et al. (Ekman et al. 2012; Grady, Thiemann, and Brockmann 2012). Therefore, we directly focused on the aroused functional connections by analyzing a wide range of different activation levels. Essentially, rather than recruiting node-consequent edges, we directly collected edge activities by specific criteria. Namely, we divided the temporal horizon of the events in two succeeding functional phases contriving a condition for the elicitation of the edges. At first, each model network was exerted 100 times. In the first trial, the algorithm parted edges such that where EBC was smaller or equal to its first distribution percentile it belonged to the first group, the rest of edges to the second group. Similarly, in the second step the algorithm classified by leveraging at the second percentile and so forth until the last percentile. We investigated such dynamics for a variable size spanning 6 dyadic scales (2^5^ to 2^10^) and for each network model (higher scales were prohibitively costly in terms of computational time). Therefore, for each generated network we computed the network statistics (C and L) to track the evolution of the functional evoked network modifications.

Results from simulations were subsequently filtered to discard singularity cases where C or L was equal to 0 and to select the regions, in terms of network size and leverage range, where the hypothesis was satisfied. We called such regions as *admissible* meaning that if such a network elicits first a certain percentage (leverage point) of edges (functional graph of stage 1) then it would elicit the rest of the edges (functional graph of stage 2), we verified our hypothesis (Figure 4).

**Figure 4.**
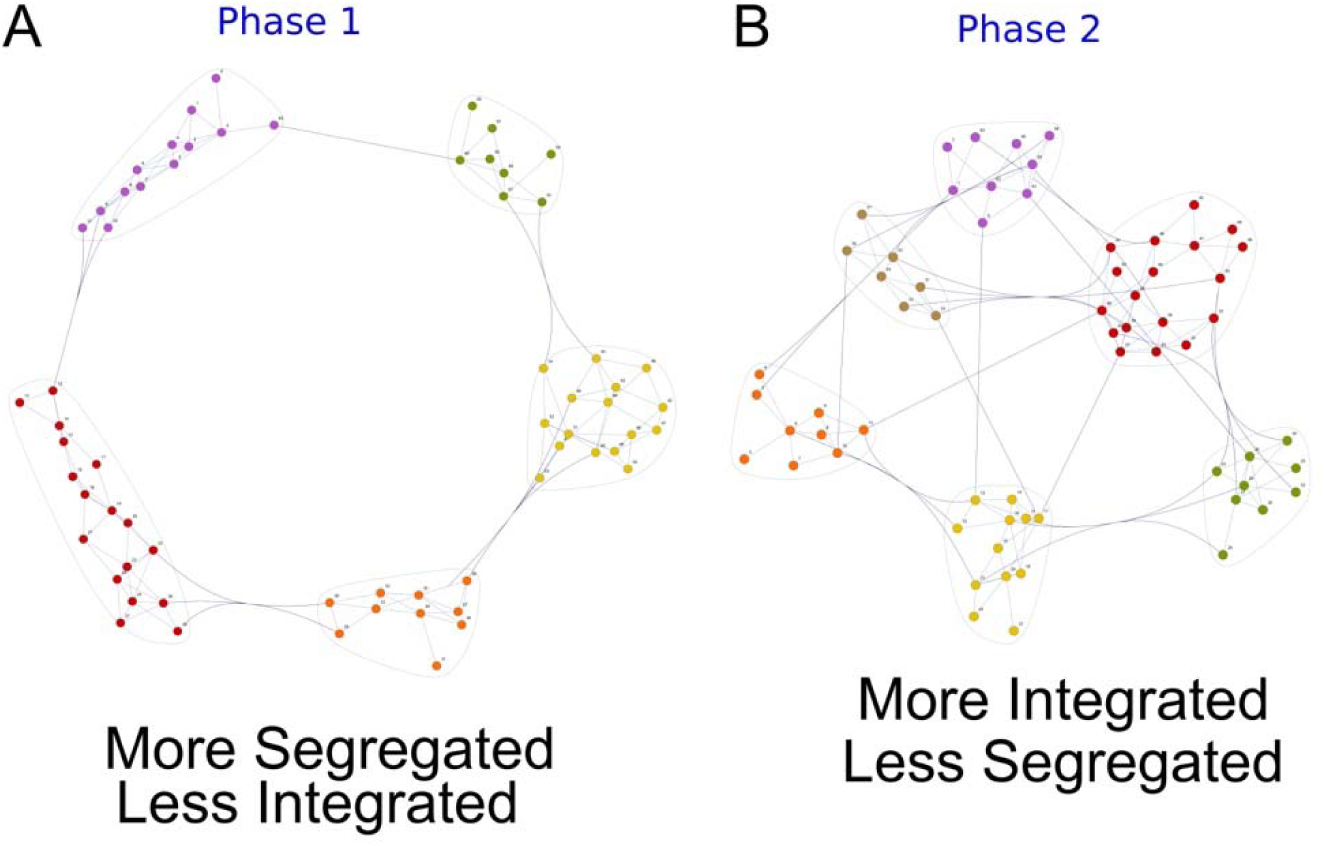
Example of the biphasic network dynamics. (A) Hypothesized “Phase 1” characterized by a functional segregation prevalence. (B) Hypothesized “Phase 2” characterized by a functional integration dominance. Both configurations were generated by simulator which used small-world networks with 64 nodes.

We found that the WS and BA models produced relevant sets of admissible regions (see Figure 5A’-D’), namely intervals of the edge leverage that verified the working hypothesis. Such regions rose with the network size in case of WS networks and shifted towards higher leverage points for BA networks. Importantly also ER models reported a considerable range of admissible regions indicating that purely random networks can be in accordance with the experimental results. But RL networks generated a quasi-empty set of admissible regions asserting that RL did not support the proposed hypothesis.

**Figure 5.**
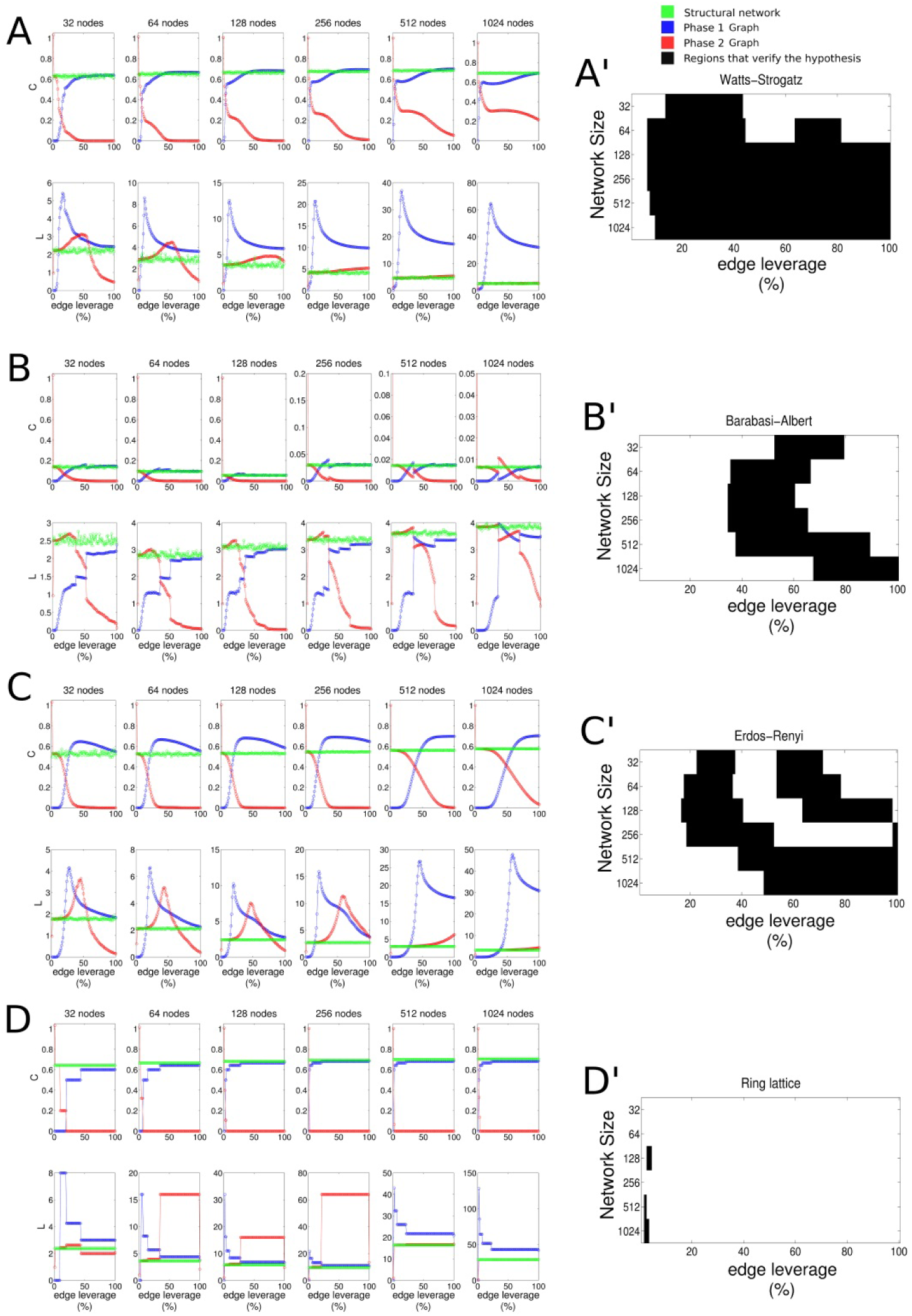
Network statistics of the network models for sizes ranging from 32 to 1024 in dyadic scales. Analyzed models were: the Watts-Strogatz (A), the Barabasi-Albert (B), the Erdos-Renyi (C) and the ring lattice (D). The first row of each subfigure indicates the Clustering Coefficient while the second one the Characteristic Path Length. Besides, subfigures A’-D’ represent the related admissible regions respectively computed for each model. More plausible models are the Watts-Strogatz and the Barabasi-Albert even though the Erdos-Renyi shows a good consistence with our hypothesis despite its much lower plausibility as brain topology.

Specifically, in WS model, when network size grew (greater or equal to 128 nodes) edge leverages nearby or greater to 10% satisfied the working hypothesis. The BA model instead was compatible with leverages in the range from ~40% to 60-65% for network of size greater or equal to 64 nodes except for the case with 1024 node wherein the admissible leverages were those from 70% or higher. The ER model was compatible in a double interval of admissible regions that were [~20%, ~40%] and [~60%, ~80%] (for network of sizes 32, 64, 128, 256), the second interval shifting to higher values according to the increasing network size. Networks of size 512 or 1024 had a single admissible region that was [40-50%, 100%]. At last, the RL model was only compatible with a very narrow range of regions and for few network sizes.

By considering the functional changes provoked by the imposed dynamics in comparison to the structural substrate, we noted that in the WS, BA and ER models the Phase 1 and Phase 2 networks largely differed from the original network (green points and excluding C and L values equal to 0), a phenomenon not observed in the RL model. A more representative example where the phenomenon is easily appreciable can be found in the movie (Movie 1) built with the C dynamics of a synthetic SWN with 128 nodes.

The evidence collected at this stage suggests a stable computational scheme for the information processing within the functional brain organization where each computation might be decomposed in sequences of two atomic and alternating steps (segregation and integration).

### Theoretical Significance

From the last section we concluded that network models, where nodes have a heavy-tailed distribution, were potentially consistent with our empirical observations from EEG activity. We further propose that these centrality distributions suggest a hierarchical processing likely divided in three layers.

A part of the experimental results showed invariances of global centralities within task trials, an important factor in the present study (Figures 6 and 7). By taking into account that the betweenness centrality (BC) in the synthetic network models used in the previous section have a similar heavy-tailed shape (Figure 8A, except for RL) of those observed in EEG functional networks (Figure 8B, for a wide range of binarization thresholds), we assumed that BC could predict the network node roles thus representing a sort of estimator of the structural-to-functional network mapping (Ekman et al. 2012; Kumar et al. 2013; Vlachos, Aertsen, and Kumar 2012; Zippo, Storchi, et al. 2013; Goh, Kahng, and Kim 2001; Zippo, Gelsomino, et al. 2013).

**Figure 6.**
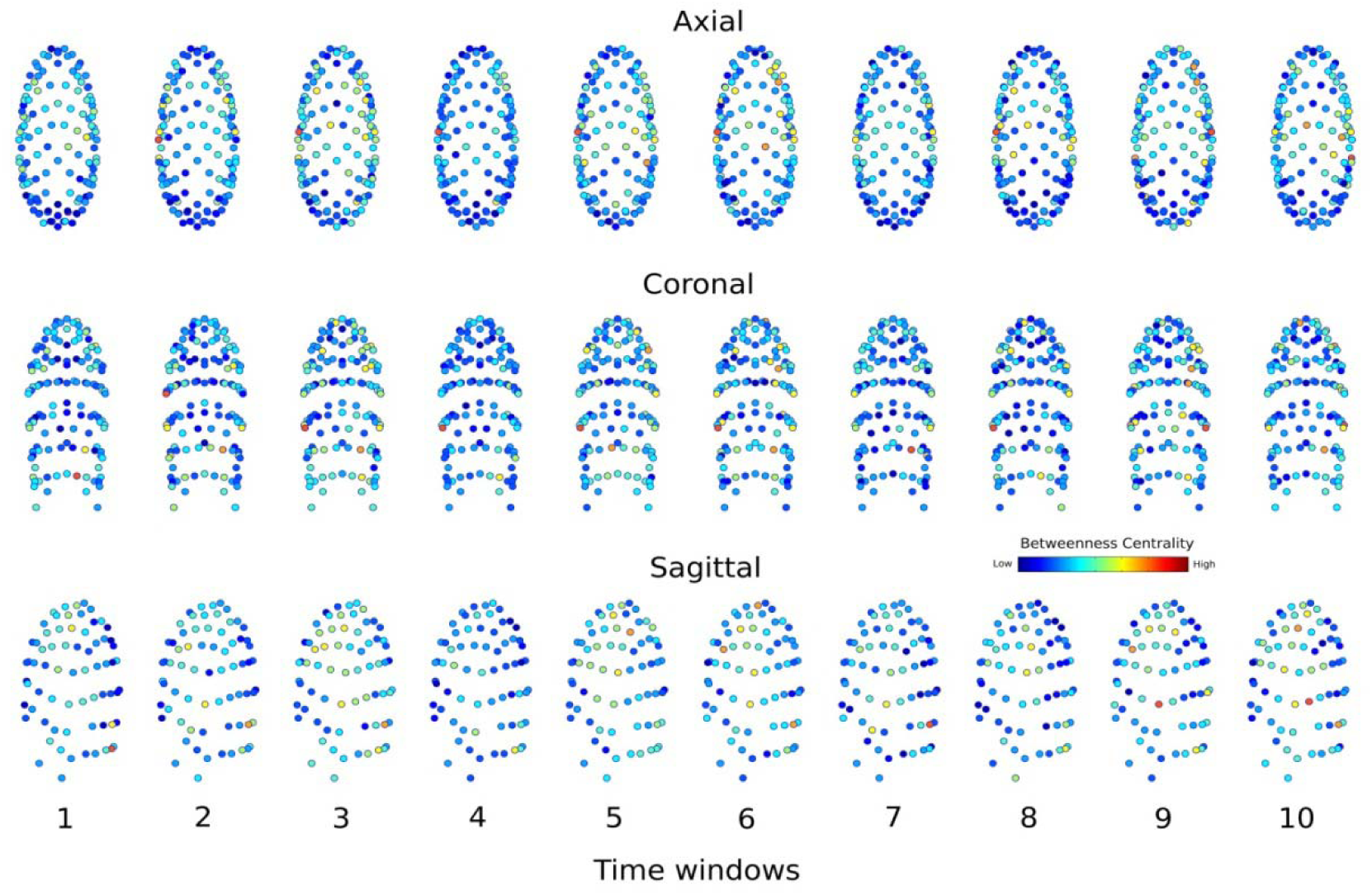
Betweenness centrality distribution over the EEG electrodes. Instance of the 2-back trial depicted in the BESA sphere space where nodes correspond to the EEG electrodes (128) and numbers (1 to 10) correspond to the time windows. Values are averaged on all trials and subjects. Node sizes are constant and node colors indicate the level of betweenness centrality.

**Figure 7.**
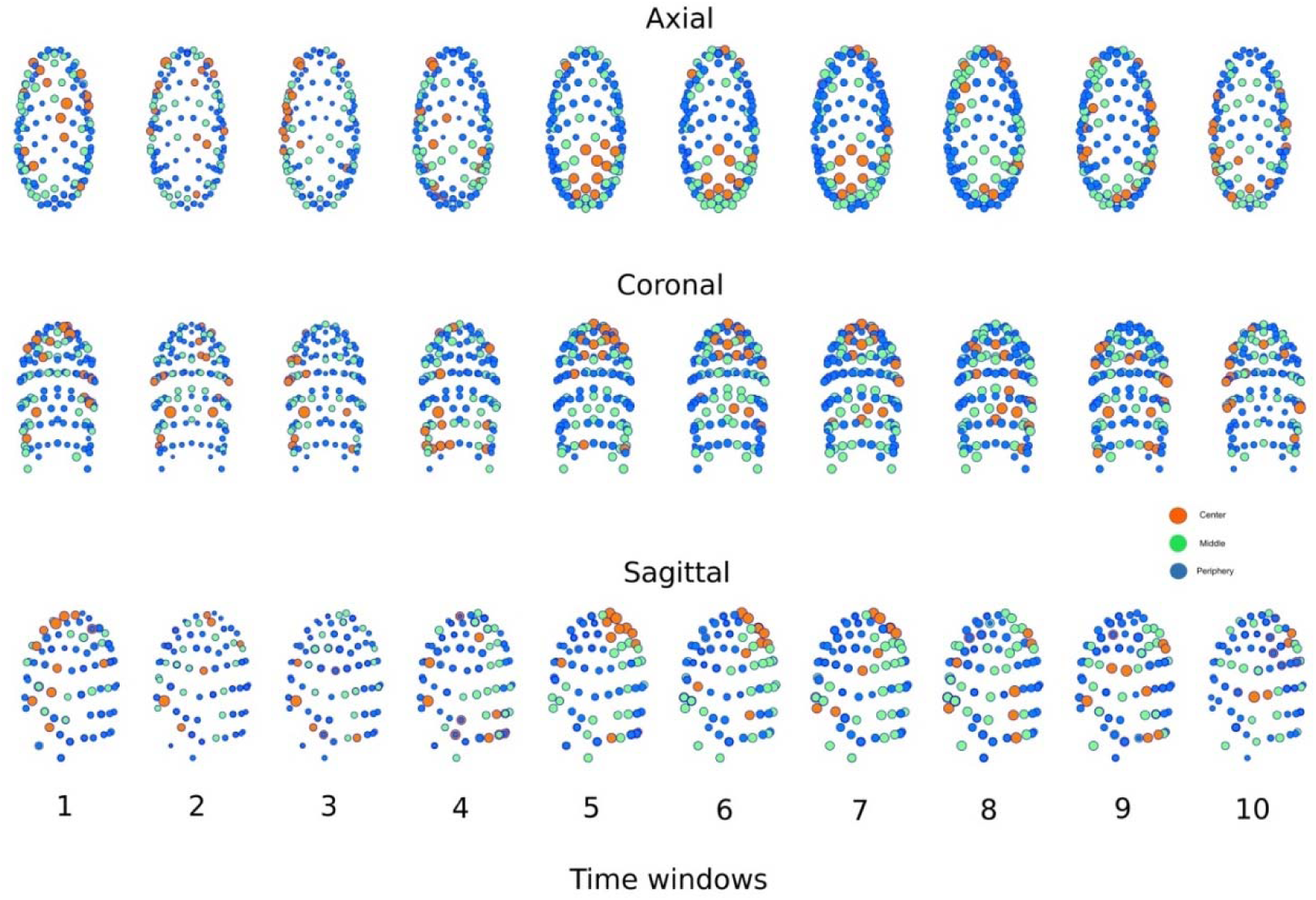
Instance of the 2-back trial depicted in the BESA sphere space where nodes correspond to the EEG electrodes (128) and numbers (1 to 10) correspond to the time windows. Node sizes are proportional to the node degree distribution and node colors indicate the level of betweenness centrality (low → network periphery, high → middle, extremely high → network center). To note that smaller nodes mean smaller node degree and, since they were tightly correlated, smaller L. In particular, the figure shows how networks evolved during a representative single trial where the node size is proportional to the node degree and the node color is referred to the BC class (red for the class iii), green for the class ii) and blue for the class i)). The windows 2, 3 and 4 highlight core activations (with nodes that switch to red and become larger) of the parietal electrodes likely recording the dorsal stream activity emergent during visual guided tasks [1,2]. This appears to represent the phase of functional segregation dominance. Subsequently, they become progressively smaller and smaller in the next windows 5, 6, 7 as potential sign of the incoming integrative processes. (Above) Axial view, (Middle) Coronal view, (Below) Sagittal view.

**Figure 8.**
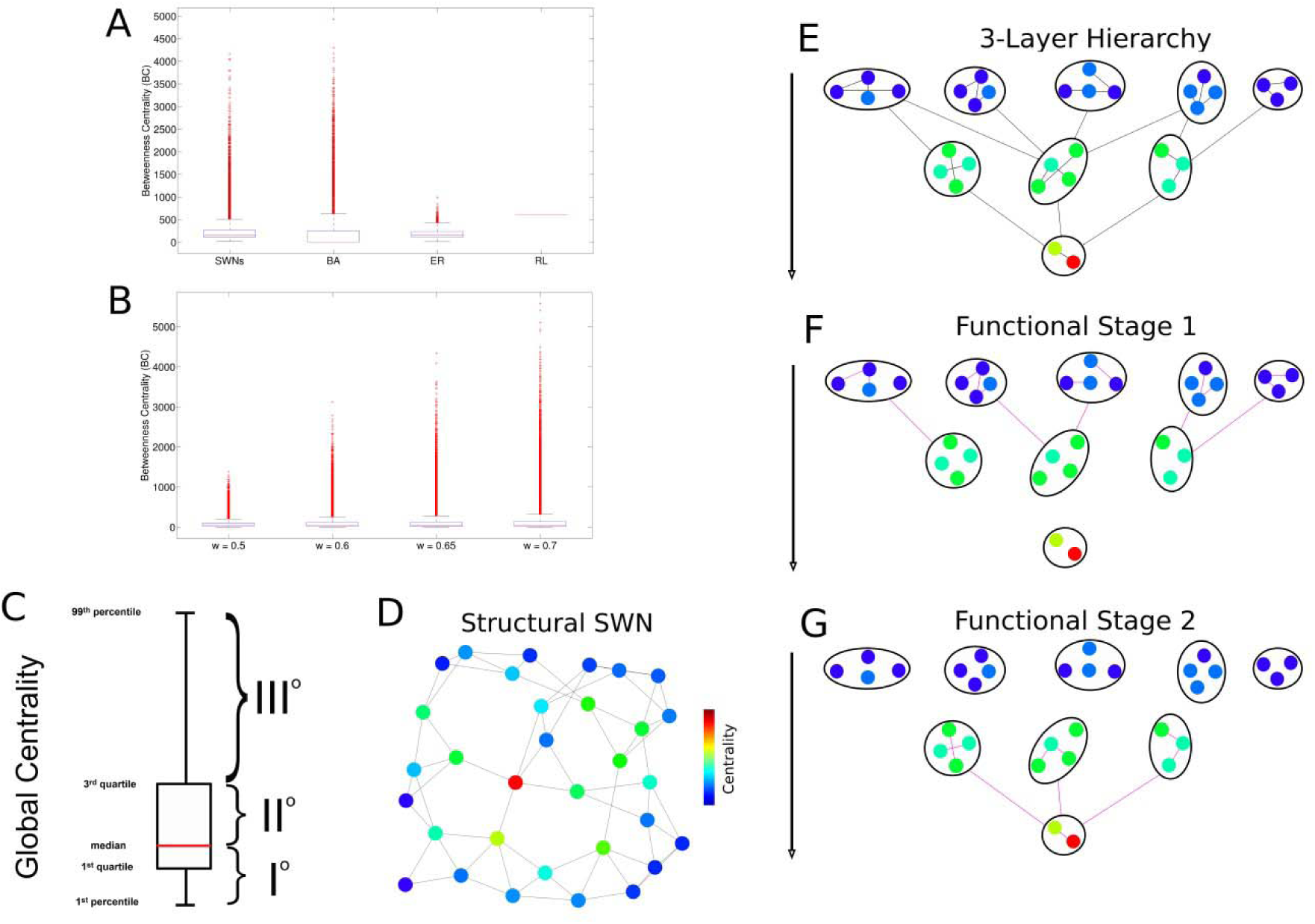
(A) Comparisons between the betweenness centrality (BC) computed on the considered network models, namely the Watts-Strogatz (SWN), the Barabasi-Albert (BA), the Erdos-Renyi (ER) and the ring lattice (RL). The SWN, BA and ER showed and heavy-tailed distribution of BC similar to those computed in EEG graphs with different thresholds. (C) The heavy-tailed shape of betweenness centrality can be reasonably parted in the classes: I) values from first to fiftieth percentiles; II) values from fifty first percentile to third quantile; III) values from seventy first percentile to ninety ninth percentile. (D) The betweenness centrality distribution on the node of a small-world network. Blue denotes nodes with low centrality, red denotes nodes with high centrality. (E) Nodes with low levels of centrality lay on the first layer of the hierarchy (different blue shades), nodes with high levels of centrality lay on the second layer of hierarchy (different green shades) and nodes with highest levels of centrality lay on the third layer (yellow and red). (F) A hypothetical stimulation of a first layer node portion activates edges (the functional connections, displayed in violet) and is named Functional Stage 1. (G) In the subsequent time step, Functional Stage 2, the activity spreads on the second and third layer nodes returning another set of activated edges.

Thus, we propose a toy network model, where nodes with low BC represent the periphery of the network and nodes with highest BC represent the core of the network, arguing that these classifications could capture the essence of the alternating phenomenon. Specifically, the toy network has a 3-layer hierarchical layout by partitioning the BC values in three arbitrary classes suggested by the BC distribution shape (see Figure 8C), with edges oriented from periphery to core (Figure 8D-E). In the ideal information flow within the toy network, a relevant part of the peripheral nodes (in layer I) are activated, then triggering a sparse activation of other layer I nodes and of a number of layer II nodes. Subsequently, activated layer II nodes similarly provoke activations of layer III nodes. Collectively, the hierarchical dynamics might be inherently reduced to two stages: activations from layer I to layer II and activations from layer II to layer III (see Figure 8F-G). In this toy network, the functional connectivity graph of each stage is acquired by inspection of the activated edges in the current stage (violet edges in Figure 8F-G). Intuitively, in the first stage the functional segregation strongly prevails on functional integration because modules activate their inner connections but remain mutually isolated (Figure 8F). Conversely, in the second stage, the functional integration dominates because, although fewer modules are active (being now first layer modules, the most conspicuous, inactive), they have gone tightly connected together.

### Discussion

In this work we investigated by EEG in healthy volunteers the global brain connectivity events during a working-memory task. We found that the temporal evolution of the involved brain network architecture follows steadily a general simple 2-step scheme wherein a surge of functional segregation flows into integration throughout the elaboration of a working-memory task. This mechanism could represent an elementary paradigm orchestrating the brain information processing in small-world networks and, hence, in effectual brain functional dynamics. To confirm this, trials where participants got wrong diverged from this rule.

### Previous Works

In these last years the study of temporal networks, namely functional networks changing their architectures in time, has progressively encompassed a widening range of disciplines. Temporal networks play an obvious critical role in the studies on brain network dynamics (Hutchison, Womelsdorf, Gati, et al. 2013; Chu et al. 2012; Sporns 2013a; Allen et al. 2014; Lefebvre 2008). Important papers have forerun crucial issues on brain circuits interpreted as temporal networks. Namely, Betzel et al. studied the repertoire of distinct states encountered by brain functional networks in EEG resting activity observing a limited set of strongly recurrent network states (Betzel et al. 2012) resembling the EEG microstates (fast and transient electrical configurations on the scalp) described elsewhere(Van de Ville, Britz, and Michel 2010). In accordance with these results, we propose here a dynamic network model capturing the early stages of a cognitive task in two connectivity states with inherent recurrences. In accordance with our results, it has been recently reported that during different types of tasks, networks showed higher degrees of integration (Crossley et al. 2013). In contrast, Kitzbichler and colleagues recently reported that global and local efficiencies of functional brain networks showed stable patterns during n-back tasks (Kitzbichler et al. 2011) and this mismatch might be ascribed to problems inherent to distortions induced by volume conductions, potentially injecting masking effects over putative neural sources.

### Brain Network Physiology

Besides these dynamic variables, our results indicate also that a hierarchical information processing could be nested into the alternating segregation and integration couples observed in trials of a working-memory task. Namely, the differences observed between 1-back and 2-back trials in terms of segregation (C) may be ascribed to the fact that a 1-back trial recruits mainly attentional processes in order to confront two successive trials, while a 2-back trial entails both attentional and control processes, the former in order to process the stream of stimuli, the latter to monitor intervening items and inhibit competing responses allowing successive integration of the information for correct response selection.

Segregation may constitute the groundings of the first stages of cognitive processing in 2-back trials when 1) a greater quantity of information needs to be held on-line in order to effectively fulfill the task goals and 2) potential sources of noise (i.e. internal or external) need to be hindered to avoid interference with task-relevant information, thus setting the premises for functional integration between salient processing elements.

Furthermore, a 2-back trial taxes cognitive processing in terms of WM load to a higher degree, which seems to translate in an increase in segregation but also modular consistency across participants in WM (Pessoa et al. 2002). Modularity is a distinctive property of a complex and efficient biological system, which tends to establish only sparse connections between sub-networks in order to scale down the propagation of noise in the system granting integration of information for demanding tasks.

According to our results, working memory can be theorized as a modular system requiring high levels of segregation in the first stages of cognitive processing both to maintain salient information no longer available and to halt interference from internal or external noise generated by competing targets. Segregation is then followed by integration in order to share and efficaciously tailor information to the specific task objectives. Wrong trials, where participants failed to get correct answers, showed very divergent network dynamics. Our synthetic network models built with the nomological scaffold as from natural data supported the prospected theory. Essentially, our results suggest that brain networks observed in the functional substrates emerge as recurrent dynamics figures virtually observable at diverse spatial and temporal scales in agreement with the findings that many brain physiology episodes are scale-free (Ekman et al. 2012). The faulty “integration-segregation” scheme in wrong trials might lead to wonder how much many brain diseases associated to cognitive impairments could harbor interfering mechanisms or exhibit weakened local dynamics to generate sound state cycling. This would also suggest that functional networks may also represent a powerful tool to discriminate normal conditions from a large repertoire of diseases.

This suggestive two-step figure doesn’t appear a standout in the physiology of living systems offering many examples of binary discrete phases such as diastoles and systoles for the hearth, inhalations and exhalations for the lung, relaxations and contractions for the peristalsis, etc. Hence, segregations and integrations for brain functional networks should not appear a remote concept nor an exceedingly reductive mechanism yet compared with the huge repertoire of states expressible by the human brain.

### Simulation interpretations

Several synthetic network models could produce the functional dynamics observed in our experiments. The simulation endorsed the possibility that the mechanisms observed in EEG sessions are due to the inherent topological brain organization which displays specialized modules able to convey processed information in fast communicating central modules of interconnected hubs. Simulated networks preferred an edge-centric perspective of the network dynamics because classical studies of neuronal network dynamics which use node behaviors (e.g. integrate and fire, Izhikevich, Hodgkin-Huxley models) are dramatically affected by the choice of model and parameters.

Although we hypothesized and verified that small-world and core-periphery networks were consistent with experimental data, we unexpectedly found that also random networks could support the observed topological phenomena. Indeed, although brain networks are far from being random, random networks have a heavy-tailed distribution of the node (or edge) centrality and we identified this property as the main cause for the detected topological phenomenon.

The *gedankenexperiment* further suggested that the observed 2-step frame could be the result of a stable hierarchical information processing layout, organized in three layers, periphery, median and core nodes, in networks with a modules-and-hubs organization. Such an organization suggests a specific computational workflow where parallel computations in segregated modules (with low centrality) spread activity to the second layer of the hierarchy (segregation stage). The activated second layer nodes (hubs) inject the obtained computed information into the last most central nodes, which reside in the third hierarchy layer (integration stage). Although the hierarchical layout in our network model based on the node centrality was postulated, the brain hierarchical organization and the hierarchical information processing in neural circuits have been largely reported. Although the bottom-up hierarchical layout in our network model based on the node centrality was postulated, the brain hierarchical organization and the hierarchical information processing in neural circuits have been largely reported (Zippo, Storchi, et al. 2013; Zippo, Gelsomino, et al. 2013; Riesenhuber and Poggio 1999; Meunier, Lambiotte, and Bullmore 2010).

### Limitations and Conclusions

A note has to be spent on earlier events that may generate, modulate or influence the double step of segregation and integration in these memory tasks. Precocious signs of segregation are detectable at 300 ms from the start of the task and flourish throughout the time window up to 700 ms. Timings appear consistently overlapping with P300 waves, at least with later component of P300, the so called the P3b associated to information processing (Squires, Squires, and Hillyard 1975). How much these multifarious components may contribute to the complex and late events of segregation and integration remains to be elucidated. As well, it remains unanswered which potential roles that precursor episodes of task error detection such as the error related negativity (Ne or ERN) and positivity (Pe), a couple of error monitoring processes, may have in interfering with the ensuing correct development of high cognitive-related processing of memorization (Falkenstein et al. 1991; Gehring et al. 1993). In the presence of Ne and Pe there could be generated destructive conditions leading to the abortion of the memorization processing. This is particularly important when considering that late P300 (a cognitive decisional ERP label) and Pe waves may represent a complex but partially overlapped neural processing with only slight temporal shifts (where Pe is present). In summary, it still remains to be clarified if in the presence of early errors, such as an incorrect motor planning, this may then drive a downstream deconstruction of the complex duet of segregation-integration and its behavioral counterpart related to memorization. Eventually, further studies are necessary to investigate the relations with the brain structural substrate. Unfortunately, no anatomical inference was possible with the available experimental setup.

In conclusion, the results in this study support the idea that, facing space-time limited context tasks, the human brain functional networks may work in accordance with two-step rules. Such rules could, further, be a natural consequence of the hierarchical information workflow of those networks. Therefore, the fluctuation repertories observed in brain functional networks might be elucidated by equivalent network mechanisms that would expand our comprehension of human brain network dynamics.

### Information Sharing Statement

All data are available upon request due to privacy restriction on human subjects and only for scientific research scopes. Requests should be submitted to the corresponding author. The entire computational framework is freely available, and it is able to reproduce data and figures (https://sites.google.com/site/antoniogiulianozippo/codes, section “Information Flow in Brain Network Topologies”).

## Acknowledgments

This work was partially supported by PON project 01-01297 (Virtualab). The funders had no role in study design, data collection and analysis, decision to publish, or preparation of the manuscript. The authors have declared that no competing interests exist. We wish to thank Ms. Chiara D’Aversa, Mr. Pieter Van Duin and Mr. Gian Carlo Caramenti for their helpful contributions.

---

**Movie 1.** The movie shows a network flow, in terms of functional connection graphs, simulated by a small-world network with 128 nodes tracking the evolution of the clustering coefficient (C). Remarkably, it is clearly evident the oscillation behavior of C that because it was tightly correlated with L visually demonstrate that such networks underwent to two topologically and orthogonal phase likely supporting the information processing demand.

## Contributions

A.G.Z. conceive the rational of the work, performed the experiments and analyzed the data. A. G. Z. wrote the manuscript together with G.E.B, P.A.D. and I.C. A.G.Z. designed the study. All of the authors made important suggestions to the manuscript and reviewed and approved the manuscript.

## Corresponding author

Correspondence to Antonio Giuliano Zippo ().

## References

Allen, Elena A., Eswar Damaraju, Sergey M. Plis, Erik B. Erhardt, Tom Eichele, and Vince D. Calhoun. 2014. “Tracking Whole-Brain Connectivity Dynamics in the Resting State.” Cerebral Cortex 24 (3): 663–76. doi:10.1093/cercor/bhs352.

Baddeley, Alan. 2003. “Working Memory: Looking Back and Looking Forward.” Nature Reviews. Neuroscience 4 (10). England: 829–39. doi:10.1038/nrn1201.

Baddeley, Alan, and Barbara A Wilson. 2002. “Prose Recall and Amnesia: Implications for the Structure of Working Memory.” Neuropsychologia 40 (10). England: 1737–43.

Betzel, Richard F, Molly a Erickson, Malene Abell, Brian F O’Donnell, William P Hetrick, and Olaf Sporns. 2012. “Synchronization Dynamics and Evidence for a Repertoire of Network States in Resting EEG.” Frontiers in Computational Neuroscience 6 (September): 74. doi:10.3389/fncom.2012.00074.

Blinowska, Katarzyna J, Maciej Kamiński, Aneta Brzezicka, and Jan Kamiński. 2013. “Application of Directed Transfer Function and Network Formalism for the Assessment of Functional Connectivity in Working Memory Task.” Philosophical Transactions of the Royal Society of London A: Mathematical, Physical and Engineering Sciences 371 (1997). The Royal Society: 20110614.

Borgatti, Stephen P. 2005. “Centrality and Network Flow.” Social Networks 27 (1): 55–71. doi:10.1016/j.socnet.2004.11.008.

Bullmore, Ed, and Olaf Sporns. 2009. “Complex Brain Networks: Graph Theoretical Analysis of Structural and Functional Systems.” Nature Reviews. Neuroscience 10 (3). England: 186–98. doi:10.1038/nrn2575.

Bullmore, Ed, and Olaf Sporns. 2012. “The Economy of Brain Network Organization.” Nature Reviews. Neuroscience 13 (5). England: 336–49. doi:10.1038/nrn3214.

Chu, Catherine J, Mark a Kramer, Jay Pathmanathan, Matt T Bianchi, M Brandon Westover, Lauren Wizon, and Sydney S Cash. 2012. “Emergence of Stable Functional Networks in Long-Term Human Electroencephalography.” The Journal of Neuroscience?: The Official Journal of the Society for Neuroscience 32 (8): 2703–13. doi:10.1523/JNEUROSCI.5669-11.2012.

Crossley, Nicolas a, Andrea Mechelli, Petra E Vértes, Toby T Winton-Brown, Ameera X Patel, Cedric E Ginestet, Philip McGuire, and Edward T Bullmore. 2013. “Cognitive Relevance of the Community Structure of the Human Brain Functional Coactivation Network.” Proceedings of the National Academy of Sciences of the United States of America 110 (28): 11583–88. doi:10.1073/pnas.1220826110.

Delorme, Arnaud, and Scott Makeig. 2004. “EEGLAB: An Open Source Toolbox for Analysis of Single-Trial EEG Dynamics Including Independent Component Analysis.” Journal of Neuroscience Methods 134 (1). Netherlands: 9–21. doi:10.1016/j.jneumeth.2003.10.009.

Ekman, Matthias, Jan Derrfuss, Marc Tittgemeyer, and Christian J Fiebach. 2012. “Predicting Errors from Recon Fi Guration Patterns in Human Brain Networks.” Proc Natl Acad Sci USA 109: 16714–19. doi:10.1073/pnas.1207523109//DCSupplemental.www.pnas.org/cgi/doi/10.1073/pnas.1207523109.

Falkenstein, M, J Hohnsbein, J Hoormann, and L Blanke. 1991. “Effects of Crossmodal Divided Attention on Late ERP Components. II. Error Processing in Choice Reaction Tasks.” Electroencephalography and Clinical Neurophysiology 78 (6): 447–55. doi:http://dx.doi.org/10.1016/0013-4694(91)90062-9.

Freeman, Linton C. 1977. “A Set of Measures of Centrality Based on Betweenness.” Sociometry 40 (1). [American Sociological Association, Sage Publications, Inc.]: 35–41. doi:10.2307/3033543.

Gehring, William J, Brian Goss, Michael G H Coles, David E Meyer, and Emanuel Donchin. 1993. “A Neural System for Error Detection and Compensation.” Psychological Science 4 (6): 385–90. doi:10.1111/j.1467-9280.1993.tb00586.x.

Goh, K I, B Kahng, and D Kim. 2001. “Universal Behavior of Load Distribution in Scale-Free Networks.” Physical Review Letters 87 (27 Pt 1). American Physical Society: 278701. doi:10.1103/PhysRevLett.87.278701.

Gordon, S. M., P. J. Franaszczuk, W. D. Hairston, M. Vindiola, and K. McDowell. 2013. “Comparing Parametric and Nonparametric Methods for Detecting Phase Synchronization in EEG.” Journal of Neuroscience Methods 212 (2). Elsevier B.V.: 247–58. doi:10.1016/j.jneumeth.2012.10.002.

Gould, P. R. 2016. “On the Geographical Interpretation of Eigenvalues” 44 (2): 15–41.

Grady, Daniel, Christian Thiemann, and Dirk Brockmann. 2012. “Robust Classification of Salient Links in Complex Networks.” Nature Communications 3 (May). Nature Publishing Group: 864. doi:10.1038/ncomms1847.

Humphries, Mark D, and Kevin Gurney. 2008. “Network ‘Small-World-Ness’: a Quantitative Method for Determining Canonical Network Equivalence.” PloS One 3 (4). United States: e0002051. doi:10.1371/journal.pone.0002051.

Hutchison, R. Matthew, Thilo Womelsdorf, Elena A. Allen, Peter A. Bandettini, Vince D. Calhoun, Maurizio Corbetta, Stefania Della Penna, et al. 2013. “Dynamic Functional Connectivity: Promise, Issues, and Interpretations.” NeuroImage 80. Elsevier Inc.: 360–78. doi:10.1016/j.neuroimage.2013.05.079.

Hutchison, R. Matthew, Thilo Womelsdorf, Joseph S. Gati, Stefan Everling, and Ravi S. Menon. 2013. “Resting-State Networks Show Dynamic Functional Connectivity in Awake Humans and Anesthetized Macaques.” Human Brain Mapping 34 (9): 2154–77. doi:10.1002/hbm.22058.

Jaeggi, Susanne M, Ria Seewer, Arto C Nirkko, Doris Eckstein, Gerhard Schroth, Rudolf Groner, and Klemens Gutbrod. 2003. “Does Excessive Memory Load Attenuate Activation in the Prefrontal Cortex? Load-Dependent Processing in Single and Dual Tasks: Functional Magnetic Resonance Imaging Study.” NeuroImage 19 (2): 210–25. doi:10.1016/S1053-8119(03)00098-3.

Kitzbichler, Manfred G, Richard N a Henson, Marie L Smith, Pradeep J Nathan, and Edward T Bullmore. 2011. “Cognitive Effort Drives Workspace Configuration of Human Brain Functional Networks.” The Journal of Neuroscience: The Official Journal of the Society for Neuroscience 31 (22): 8259–70. doi:10.1523/JNEUROSCI.0440-11.2011.

Kumar, Arvind, Ioannis Vlachos, Ad Aertsen, and Clemens Boucsein. 2013. “Challenges of Understanding Brain Function by Selective Modulation of Neuronal Subpopulations.” Trends in Neurosciences 36 (10). Elsevier Ltd: 579–86. doi:10.1016/j.tins.2013.06.005.

Lefebvre, Vincent D Blondel and Jean-Loup Guillaume and Renaud Lambiotte and Etienne. 2008. “Fast Unfolding of Communities in Large Networks.” Journal of Statistical Mechanics: Theory and Experiment 2008 (10): P10008. http://stacks.iop.org/1742-5468/2008/i=10/a=P10008.

Meunier, David, Renaud Lambiotte, and Edward T Bullmore. 2010. “Modular and Hierarchically Modular Organization of Brain Networks.” Frontiers in Neuroscience 4 (December): 200. doi:10.3389/fnins.2010.00200.

Newman, M E J. 2006. “Modularity and Community Structure in Networks.” Proceedings of the National Academy of Sciences of the United States of America 103 (23): 8577–82. doi:10.1073/pnas.0601602103.

Oberauer, K, and H M Suss. 2000. “Working Memory and Interference: A Comment on Jenkins, Myerson, Hale, and Fry (1999).” Psychonomic Bulletin & Review 7 (4). United States: 727–40.

Oberauer, Klaus. 2003. “Selective Attention to Elements in Working Memory.” Experimental Psychology 50 (4). Germany: 257–69. doi:10.1026//1618-3169.50.4.257.

Ortiz, Erick, Krunoslav Stingl, Jana Munssinger, Christoph Braun, Hubert Preissl, and Paolo Belardinelli. 2012. “Weighted Phase Lag Index and Graph Analysis: Preliminary Investigation of Functional Connectivity during Resting State in Children.” Computational and Mathematical Methods in Medicine 2012. United States: 186353. doi:10.1155/2012/186353.

Owen, Adrian M, Kathryn M McMillan, Angela R Laird, and Ed Bullmore. 2005. “N-Back Working Memory Paradigm: A Meta-Analysis of Normative Functional Neuroimaging Studies.” Human Brain Mapping 25 (1). United States: 46–59. doi:10.1002/hbm.20131.

Palva, J Matias, Alexander Zhigalov, Jonni Hirvonen, Onerva Korhonen, Klaus Linkenkaer-Hansen, and Satu Palva. 2013. “Neuronal Long-Range Temporal Correlations and Avalanche Dynamics Are Correlated with Behavioral Scaling Laws.” Proceedings of the National Academy of Sciences of the United States of America 110 (9): 3585–90. doi:10.1073/pnas.1216855110.

Park, Hae-Jeong, and Karl Friston. 2013. “Structural and Functional Brain Networks: From Connections to Cognition.” Science (New York, N.Y.) 342 (6158): 1238411. doi:10.1126/science.1238411.

Pessoa, Luiz, Eva Gutierrez, Peter Bandettini, and Leslie Ungerleider. 2002. “Neural Correlates of Visual Working Memory: fMRI Amplitude Predicts Task Performance.” Neuron 35 (5). United States: 975–87.

Riesenhuber, M, and T Poggio. 1999. “Hierarchical Models of Object Recognition in Cortex.” Nature Neuroscience 2 (11): 1019–25. doi:10.1038/14819.

Rubinov, Mikail, and Olaf Sporns. 2010. “Complex Network Measures of Brain Connectivity: Uses and Interpretations.” NeuroImage 52 (3). Elsevier Inc.: 1059–69. doi:10.1016/j.neuroimage.2009.10.003.

Rubinov, Mikail, and Olaf Sporns. 2011. “Weight-Conserving Characterization of Complex Functional Brain Networks.” NeuroImage 56 (4). Elsevier Inc.: 2068–79. doi:10.1016/j.neuroimage.2011.03.069.

Senden, Mario, Gustavo Deco, Marcel A. De Reus, Rainer Goebel, and Martijn P. Van Den Heuvel. 2014. “Rich Club Organization Supports a Diverse Set of Functional Network Configurations.” NeuroImage 96. Elsevier Inc.: 174–82. doi:10.1016/j.neuroimage.2014.03.066.

Sporns, Olaf. 2013a. “Network Attributes for Segregation and Integration in the Human Brain.” Current Opinion in Neurobiology 23 (2). Elsevier Ltd: 162–71. doi:10.1016/j.conb.2012.11.015.

Sporns, Olaf. 2013b. “Making Sense of Brain Network Data.” Nature Methods 10 (6). United States: 491–93. doi:10.1038/nmeth.2485.

Squires, Nancy K, Kenneth C Squires, and Steven A Hillyard. 1975. “Two Varieties of Long-Latency Positive Waves Evoked by Unpredictable Auditory Stimuli in Man.” Electroencephalography and Clinical Neurophysiology 38 (4): 387–401. doi:http://dx.doi.org/10.1016/0013-4694(75)90263-1.

Stam, Cornelis J, Guido Nolte, and Andreas Daffertshofer. 2007. “Phase Lag Index: Assessment of Functional Connectivity from Multi Channel EEG and MEG with Diminished Bias from Common Sources.” Human Brain Mapping 28 (11): 1178–93. doi:10.1002/hbm.20346.

Telesford, Qawi K, Karen E Joyce, Satoru Hayasaka, Jonathan H Burdette, and Paul J Laurienti. 2011. “The Ubiquity of Small-World Networks” 1 (5). doi:10.1089/brain.2011.0038.

Tononi, G, G M Edelman, and O Sporns. 1998. “Complexity and Coherency: Integrating Information in the Brain.” Trends in Cognitive Sciences 2 (12): 474–84. http://www.ncbi.nlm.nih.gov/pubmed/21227298.

Van de Ville, Dimitri, Juliane Britz, and Christoph M Michel. 2010. “EEG Microstate Sequences in Healthy Humans at Rest Reveal Scale-Free Dynamics.” Proceedings of the National Academy of Sciences of the United States of America 107 (42): 18179–84. doi:10.1073/pnas.1007841107.

van den Heuvel, M P, C J Stam, M Boersma, and H E Hulshoff Pol. 2008. “Small-World and Scale-Free Organization of Voxel-Based Resting-State Functional Connectivity in the Human Brain.” NeuroImage 43 (3): 528–39. doi:10.1016/j.neuroimage.2008.08.010.

van den Heuvel, Martijn P, and Olaf Sporns. 2011. “Rich-Club Organization of the Human Connectome.” The Journal of Neuroscience: The Official Journal of the Society for Neuroscience 31 (44): 15775–86. doi:10.1523/JNEUROSCI.3539-11.2011.

Vinck, Martin, Robert Oostenveld, Marijn Van Wingerden, Franscesco Battaglia, and Cyriel M A Pennartz. 2011. “An Improved Index of Phase-Synchronization for Electrophysiological Data in the Presence of Volume-Conduction, Noise and Sample-Size Bias.” NeuroImage 55 (4). Elsevier Inc.: 1548–65. doi:10.1016/j.neuroimage.2011.01.055.

Vinck, Martin, Robert Oostenveld, Marijn van Wingerden, Franscesco Battaglia, and Cyriel M a Pennartz. 2011. “An Improved Index of Phase-Synchronization for Electrophysiological Data in the Presence of Volume-Conduction, Noise and Sample-Size Bias.” NeuroImage 55 (4). Elsevier Inc.: 1548–65. doi:10.1016/j.neuroimage.2011.01.055.

Vlachos, Ioannis, Ad Aertsen, and Arvind Kumar. 2012. “Beyond Statistical Significance: Implications of Network Structure on Neuronal Activity.” PLoS Computational Biology 8 (1): e1002311. doi:10.1371/journal.pcbi.1002311.

Watts, D J, and S H Strogatz. 1998. “Collective Dynamics of‘Small-World’ Networks.” Nature 393 (6684): 440–42. doi:10.1038/30918.

Whitlow, Christopher T, Ramon Casanova, and Joseph A Maldjian. 2011. “Effect of Resting-State Functional MR Imaging Duration on Stability of Graph Theory Metrics of Brain Network Methods: Results:” 259 (2). doi:10.1148/radiol.11101708/-/DC1.

Zippo, Antonio G, Giuliana Gelsomino, Pieter Van Duin, Sara Nencini, Gian Carlo Caramenti, Maurizio Valente, and Gabriele E M Biella. 2013. “Small-World Networks in Neuronal Populations: A Computational Perspective.” Neural Networks: The Official Journal of the International Neural Network Society 44 (August): 143–56. doi:10.1016/j.neunet.2013.04.003.

Zippo, Antonio G, Riccardo Storchi, Sara Nencini, Gian Carlo Caramenti, Maurizio Valente, and Gabriele Eliseo M Biella. 2013. “Neuronal Functional Connection Graphs among Multiple Areas of the Rat Somatosensory System during Spontaneous and Evoked Activities.” PLoS Computational Biology 9 (6): e1003104. doi:10.1371/journal.pcbi.1003104.

